# Audited vibe coding suggests partial fetal-like convergence of tumor proteomes

**DOI:** 10.64898/2026.08.26.745609

**Authors:** Jesse G. Meyer

**Affiliations:** Department of Computational Biomedicine, Cedars-Sinai Medical Center, Los Angeles, California, USA

## Abstract

The balance between how much human tumors recapitulate fetal tissue programs versus lose adult tissue identity remains unresolved. I used audited vibe coding, a human-mediated, cross-model critique-and-refinement workflow, to re-analyze a public pan-cancer proteomic atlas. A primary large language model wrote and executed the analysis under scientific direction, while a separate model family audited the code, outputs and claims; findings were returned for correction across seven versioned releases. Among 229 tumor-adjacent pairs in seven organs, tumor-minus-adjacent proteomic change partially aligned with reverse fetal-to-adult maturation (organ-balanced cosine, 0.240; 95% interval, 0.138 to 0.335), with positive alignment in 189 of 229 patients (82.5%). The organ-balanced projection coefficient was 0.195 (95% interval, 0.069 to 0.244), indicating movement along only part of the developmental distance. Although reverse maturation overlapped adult-identity loss, a positive developmental component remained after identity loss entered first (0.203; 95% interval, 0.129 to 0.239). Suppression of adult-high proteins contributed to more positive alignment than reactivation of fetal-high proteins. The vibe coding audits identified substantive defects. A common-mask correction reduced the matched-organ advantage from 0.074 to 0.059; a missing-value correction barely changed aggregate geometry but replaced 5 of the top 40 liver contributors; and coupled resampling repaired uncertainty accounting without changing patient scores. As with any single report, the “vibe reanalysis” biological results are candidate discoveries pending independent replication. The workflow is a single feasibility case, not a reliability benchmark, and shows how conversationally generated analysis can be made more inspectable when model-written code is treated as untrusted, versioned and subject to separate-model critique and executable checks.

## Introduction

Tumors can reactivate embryonic, fetal and stemness-associated programs, but apparent fetal similarity has more than one interpretation.^1-6^ Tumor proteomic change might reverse organ maturation, merely erode adult tissue identity or reflect a shared cancer program unrelated to development.^7,8^ Distinguishing these possibilities requires paired tumor-adjacent measurements and explicitly separated reference directions rather than a pooled fetal-tumor-adult trajectory.

The TPHP human proteome atlas provides fetal, adult-normal and paired tumor-adjacent measurements across many tissues, creating an opportunity to test this question.^9^ Like other public omics atlases,^10-13^ it does not provide every bespoke comparison that later investigators may want. A targeted re-analysis still requires custom parsing, cohort construction, geometry, uncertainty estimation, figures and claim tracking.

’Vibe coding’ is the colloquial practice of directing a large language model in natural language while allowing it to author most or all of the implementation.^14^ It can move a domain scientist rapidly from a biological question to executable analysis without manually writing every line. Here, that conversational path was embedded in a separate-model audit loop and preserved across versioned releases (**Fig. 1a,b**). For science, the central risk is that fluent code and plausible figures may conceal a broken connection between intent, implementation and claim. Benchmarks and real-world software studies likewise show that plausible generated code is not equivalent to correct code.^15-19^ A disclosure of AI assistance does not resolve that risk.

**Fig. 1.**
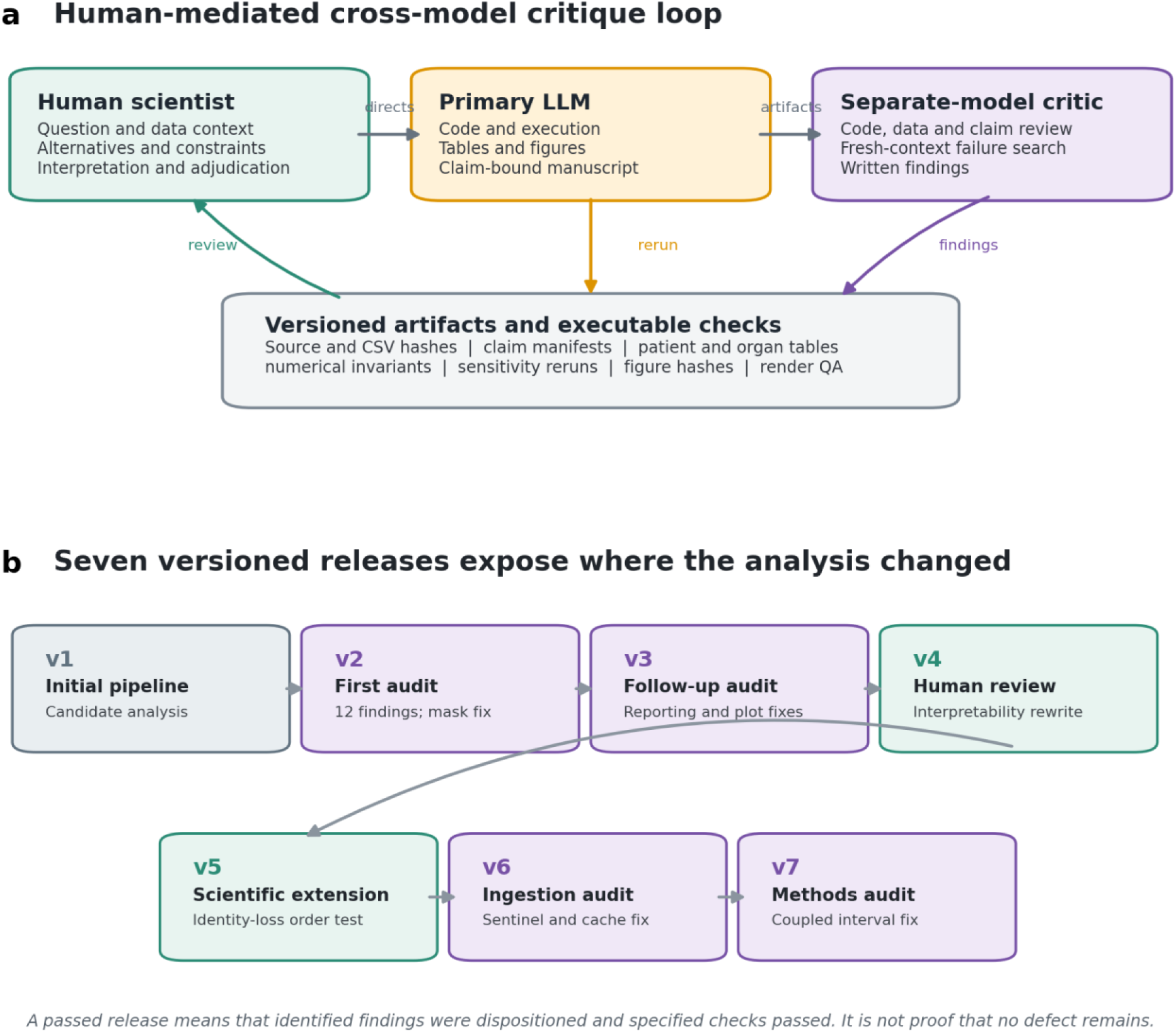
Audited vibe coding workflow and version history. a, In this human-mediated, cross-model critique-and-refinement workflow, the scientist supplied the broad question, data context, alternative explanations and interpretation; the primary model generated and executed code; a separate model family critiqued the resulting artifacts; findings returned to the primary chat; and machine-readable checks recorded their consequences. b, Seven releases show where human scientific steering and separate-model audits altered the analysis. A passed release denotes disposition of identified findings and completion of specified checks, not proof that the code is error-free.

Related work describes model reflection or self-refinement, in which a model critiques and revises its own output, and critic-based workflows, in which another model evaluates an output and returns feedback.^20-22^ I use ‘audited vibe coding’ as a memorable label for a human-mediated, cross-model critique-and-refinement workflow. It is not independent validation: a separate model family served as critic, while the scientist supplied the drive, biological constraints, and relayed and adjudicated findings.

Here I apply that workflow to ask whether tumor-minus-adjacent proteomic change points toward the fetal state after accounting for adult-identity loss and a shared cancer direction. The primary model generated the executable analysis, figures and manuscript; the separate model audited the code repository in comparison with the paper draft and claims; findings from each iterative audit were returned to the primary chat for correction and rerun. The study therefore has two linked outputs: a candidate biological model of partial fetal-like convergence and an inspectable case history of how cross-model critique changed the analysis.

## Results

### Audited vibe coding generated and corrected a test of fetal-like tumor change

The human scientist first supplied the PDF of the original paper describing the available data and asked what biological questions would be of interest to explore. The AI (GPT-5.6-sol with max reasoning) determined that it would be important to quantify whether the paired cancer change reverses fetal-to-adult maturation. The human scientist then supplied four workbooks consumed by the final pipeline, and asked the system to perform all the research, including figure generation and paper writing.

The primary biological endpoint was the directional alignment between each patient’s tumor-minus-adjacent proteomic change and the reverse fetal-to-adult maturation vector for that organ. Primary inference used 229 pairs in seven organs with replicated developmental donors. Projection distance, adult-identity loss, a leave-one-organ-out shared cancer axis and protein-level contributions were analyzed to distinguish partial developmental convergence from alternative explanations.

In the primary ChatGPT conversation, GPT-5.6 Sol with Max reasoning inspected the data structure, wrote Python code, ran analyses, created derived tables and five figures, and drafted the manuscript. The human scientist steered the biological choices and interrogated outputs through natural-language questions, including why cosine and projection panels differed, whether reverse maturation could be reduced to identity loss, how Gram-Schmidt affected axis overlap and how individual proteins contributed to a vector dot product. Those questions prompted clarification and additional analyses presented below.

To explore efficient verification of the first agentic system where no ground truth was available, the human scientist explored separate-model audits performed with Opus 5 in Claude Code version 2.1.226. The critic examined code, inputs, outputs, figures and manuscript claims. Findings from the serial audits were passed back into the primary ChatGPT thread; the primary model then changed code or prose, ran from source and emitted a newly named package. In this manuscript, ‘resolved’ means that each identified finding received a documented disposition and the specified release checks passed. It does not mean that no undiscovered defect remains.

The version history records both model audit and human scientific steering. Version 1 established the initial candidate analysis. Version 2 implemented a 12-item audit and reran from source. Version 3 addressed eight listed follow-up findings. Version 4 rewrote the results panel by panel after human questions exposed interpretability gaps. Version 5 added the alternative explanation that reverse maturation might simply be adult-identity loss. Version 6 corrected an ingestion sentinel and strengthened conversion provenance after another code audit. Version 7 repaired methods consistency and coupled bootstrap accounting. Earlier results were retained or changed explicitly rather than overwritten silently.

### Separate-model audits found consequential defects

The audit loop was not cosmetic. It identified failures at four distinct layers: comparison-set construction, data ingestion and provenance, claim generation, and uncertainty reporting (**Table 1**). Some findings affected only disclosure or portability, but others altered scientific outputs.

**Table 1.**
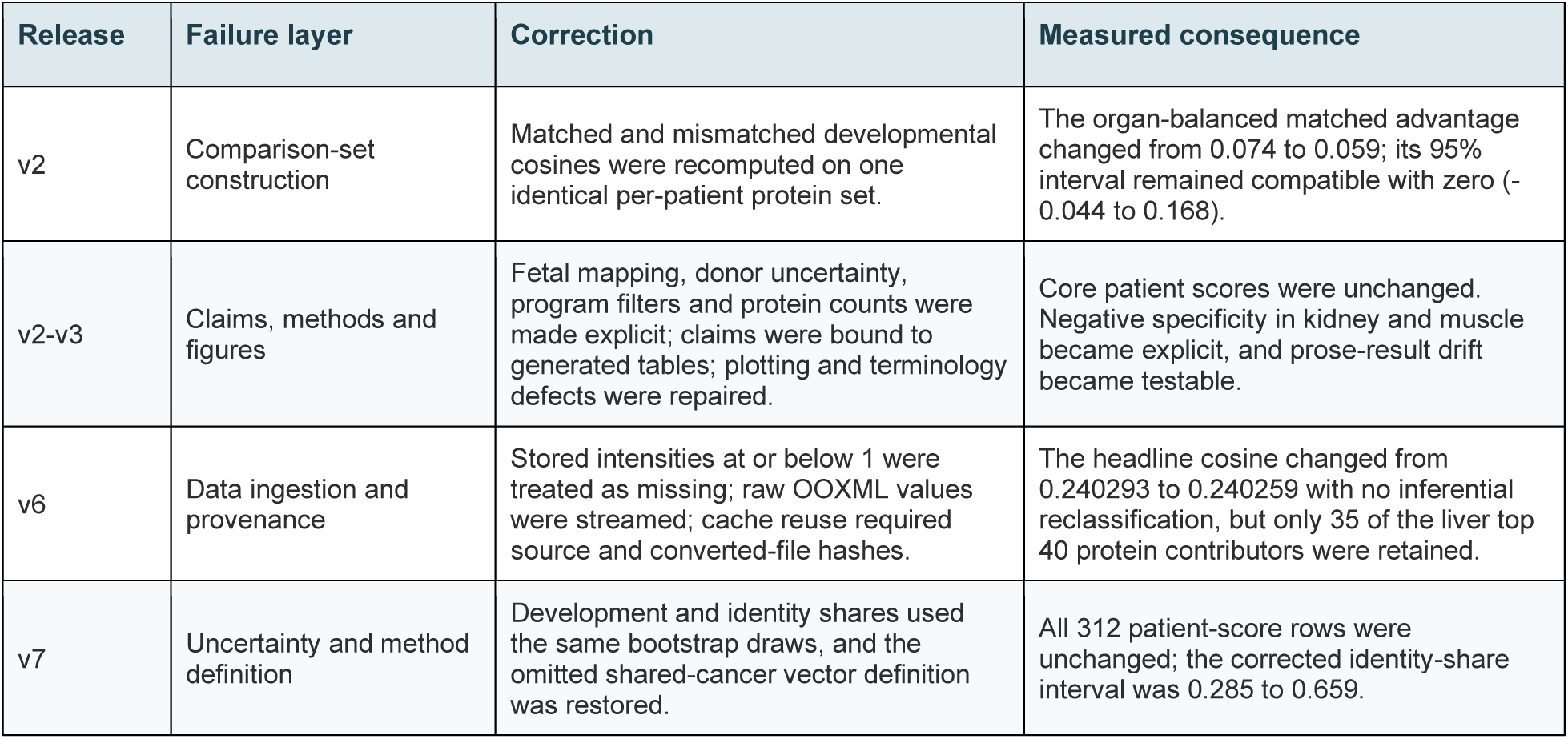
Documented findings from separate-model audits and their effects on the analysis.

The most consequential inferential correction occurred in version 2. The matched developmental cosine and the mismatched-organ cosines had not been evaluated on identical proteins within each patient. Enforcing one common comparison set reduced the organ-balanced matched-axis advantage from 0.074 to 0.059, with a 95% interval of -0.044 to 0.168. The qualitative conclusion was therefore weakened from apparent organ specificity to an interval compatible with no cross-organ advantage.

Version 6 exposed a different failure mode. Stored intensity 1 was a missing-value sentinel but had been retained as a measured abundance. The corrected matched rerun changed the headline developmental cosine only from 0.240293 to 0.240259 and changed no inferential classification. Protein ranking was less stable: only 35 of the liver top 40 contributors overlapped, so the figure and protein claims were regenerated. This distinction between a stable aggregate and a less stable mechanistic ranking would have been missed by checking only the headline result.

The final release could be rerun from the source data and included automated checks that linked each major claim and figure to its supporting results. The checks confirmed that the expected files, patients and protein values were used; that the geometric decomposition and uncertainty calculations behaved as intended; and that the manuscript and figures matched the generated results. This made known errors easier to detect and showed which conclusions changed after each correction. It did not prove that every scientific choice or line of code was correct, and both AI systems could still share blind spots.

### Geometric analysis separates developmental reversion from alternative tumor programs

The main TPHP protein-intensity matrix contains 2,856 samples and 13,609 quantified proteins.^9^ After restricting the fetal, adult-normal and paired tumor-adjacent matrices to 9,928 shared proteins and collapsing repeated acquisitions to biological units, I identified 890 complete tumor-adjacent pairs across 25 cancer types (**Fig. 2a**). The full set of pairs informed the collection of leave-one-organ-out shared-cancer references, with each target organ excluded from its own reference. Directly matched fetal, adult and cancer coverage was available for 312 pairs in 10 organs; 229 pairs in seven organs also met the replicated-donor criterion used for primary inference.

**Fig. 2.**
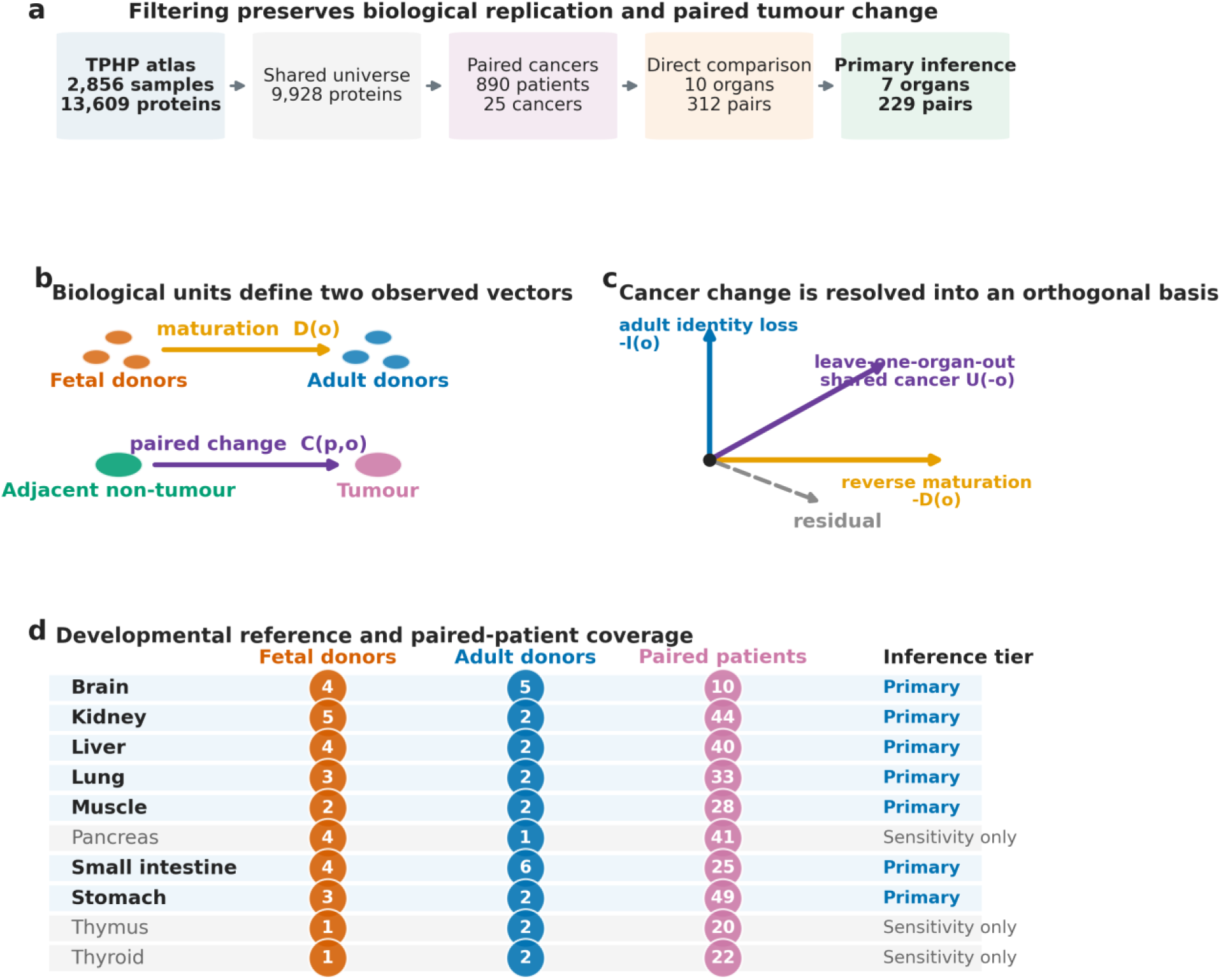
Study design and geometric decomposition of tumor-minus-adjacent proteomic change. a, Filtering from the TPHP atlas to the primary inference tier. All 890 complete pairs across 25 cancers were used to estimate leave-one-organ-out shared cancer directions; 312 pairs belonged to the 10 organs with direct fetal and adult references, and 229 pairs in seven organs met the replicated-donor criterion. b, Fetal and adult donors define maturation D(o) as adult minus fetal abundance, while paired adjacent non-tumor and tumor samples define patient cancer change C(p,o) as tumor minus adjacent abundance. Developmental reversion is -D(o). c, Cancer change is projected sequentially onto reverse maturation, adult identity loss and a leave-one-organ-out shared cancer direction; the unexplained component is the residual. Development is shown first; Fig. 4 tests the reverse order and the joint subspace. d, Biological-unit coverage by organ. Primary organs had at least two fetal and two adult donors; secondary organs were used only for sensitivity analysis.

**Figure 2b** defines the two biologically anchored vectors used to ask whether tumors move in a fetal direction. Organ maturation, D(o), is the protein-wise median adult profile minus the median fetal profile. A patient’s cancer-change vector, C(p,o), is tumor abundance minus adjacent non-tumor abundance in that same patient. Reverse maturation, -D(o), therefore points from adult toward fetal tissue. The cosine between C(p,o) and -D(o) measures their directional alignment, not the distance traveled. A positive cosine indicates that tumor change aligned with the fetal direction; zero indicates no directional alignment in the analyzed protein space; and a negative value indicates alignment with continued maturation away from the fetal reference. The distance traveled along this axis is assessed separately with the projection coefficient in **Fig. 3d**.

**Fig. 3.**
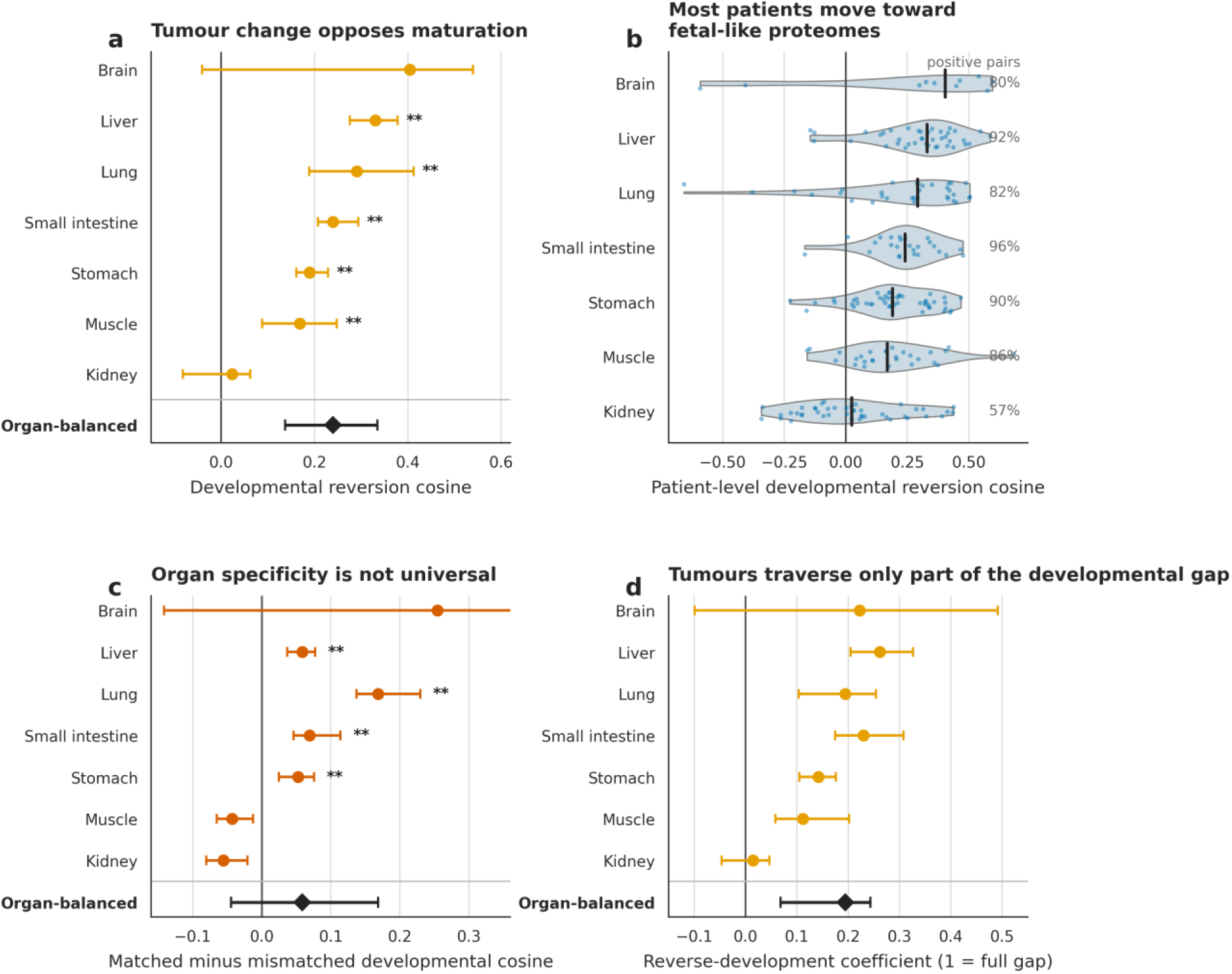
Tumor-minus-adjacent proteomic change aligns with reverse fetal-to-adult maturation. a, Median patient-level developmental reversion cosine and 95% patient-bootstrap interval for each primary organ. Positive values indicate movement toward fetal reference. The organ-balanced diamond uses an organ-patient hierarchical bootstrap. These intervals are conditional on the estimated developmental axes and do not resample fetal or adult donors. Asterisks denote Benjamini-Hochberg-adjusted sign-flip tests (*q<0.05; **q<0.01). b, Patient-level score distributions; black bars are medians and right labels are the percentage of positive pairs. c, Matched-organ cosine minus the median cosine to mismatched organ maturation axes, with all comparisons within a patient evaluated on an identical protein set (median 2,252 proteins; range, 1,154-2,278). Positive values favor the matched organ. The cross-organ interval includes zero; kidney and muscle intervals are entirely below zero. d, Projection coefficient onto reverse maturation. A value of 1 would traverse the full fetal-to-adult reference gap in reverse; 0 indicates no movement along that axis.

**Figure 2c** then projects each cancer-change vector onto a three-axis orthogonal basis constructed sequentially using the Gram-Schmidt method. The first axis is reverse maturation. The second begins with identity loss, pointing away from the target adult organ and toward the median of the other adult organs, and is stripped of its overlap with reverse maturation. The third begins with a leave-one-organ-out shared-cancer direction estimated from the other cancer organs and is stripped of its overlap with both preceding axes. The component of cancer change not captured by these three axes is retained as residual change. Because reverse maturation enters first, it receives first claim on any overlap with identity loss. This prevents double counting, but the separate development and identity scores depend on which direction enters first. **Figure 4** tests both entry orders and their order-invariant joint subspace rather than treating these operational axes as separate causal pathways.

**Fig. 4.**
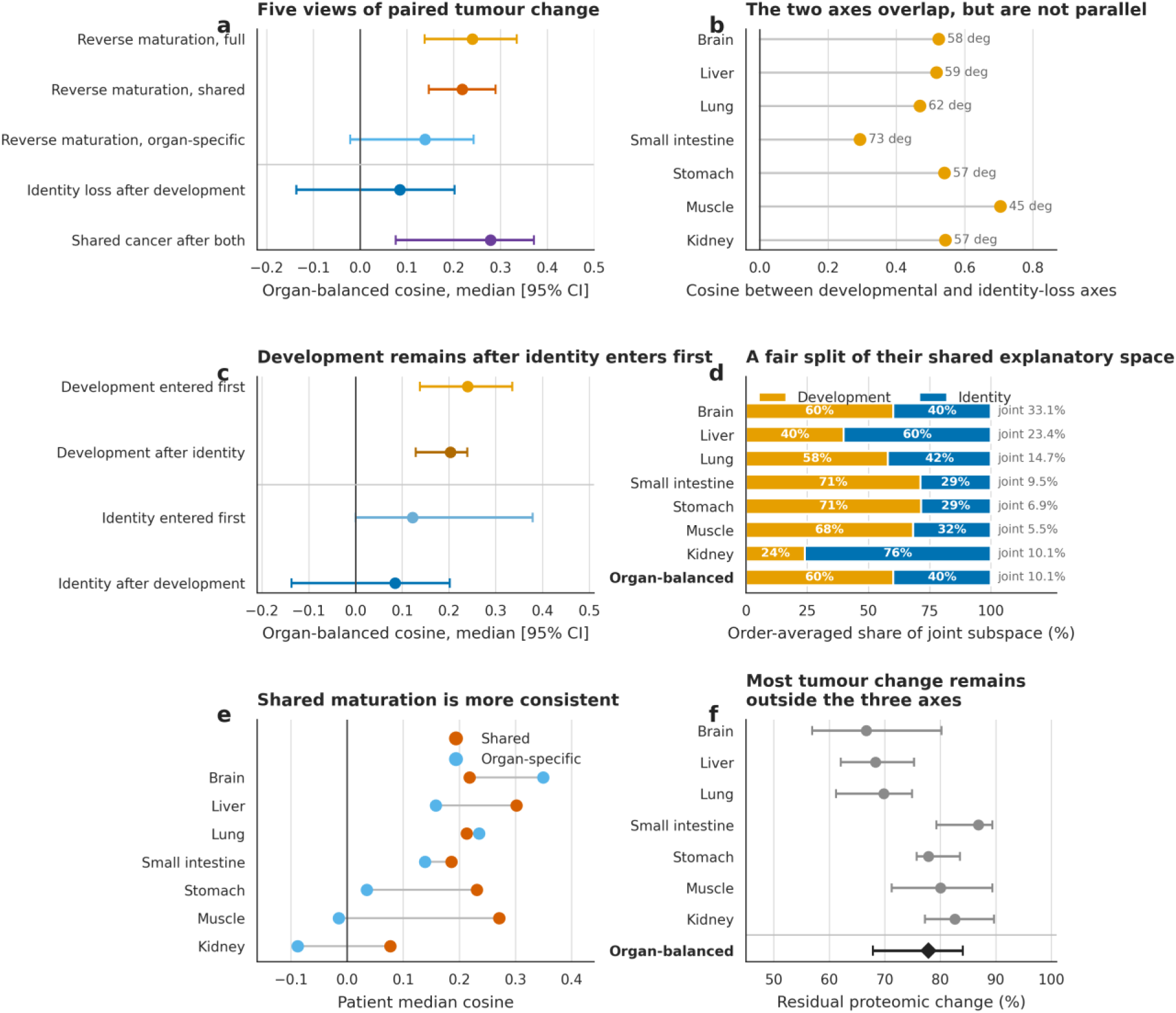
Fetal-like convergence overlaps with, but extends beyond, loss of adult tissue identity. a, Organ-balanced median cosines and 95% organ-patient hierarchical-bootstrap intervals for complete reverse maturation, its shared and organ-specific components, identity loss after development, and shared cancer after both preceding axes. ‘After’ denotes geometric residualization, not biological sequence. b, Fixed-reference cosine between reverse maturation and identity loss in each primary organ. Angle labels are arccosine transformations of the same values; no patient-bootstrap interval applies. c, Order test. Development and identity are each shown when entered first and after the other axis has been removed. d, Two-variable Shapley allocation averaged across both entry orders. Bar widths divide the joint developmental-identity subspace; labels at right give that subspace as a percentage of total paired change. The allocation is statistical, not causal. e, Paired organ medians for shared and organ-specific maturation components. f, Residual percentage after projection onto developmental, identity-loss and shared cancer axes. Bootstrap intervals are conditional on the estimated reference axes.

Finally, **Fig. 2d** makes the uneven biological support visible. Its columns give the numbers of fetal donors, adult donors and paired cancer patients, so the relatively small developmental reference cohorts cannot be mistaken for the much larger cancer cohorts. The primary tier comprised brain, kidney, liver, lung, muscle, small intestine and stomach. Pancreas, thymus and thyroid were retained only for sensitivity analyses because each had only one donor at one developmental endpoint.

### Tumor proteomic change partially aligns with reverse maturation across most organs

At the organ level, **Fig. 3a** summarizes the cosine between cancer/adjacent change and developmental reversion. Each point is an organ’s patient median, each horizontal line is its 95% patient-bootstrap interval, and the diamond gives an organ-balanced estimate, which weights organs rather than cohort sizes equally. The organ-balanced median cosine similarity was 0.240 (95% organ-patient bootstrap interval, 0.138-0.335). Five of seven organs remained significant after Benjamini-Hochberg correction of patient sign-flip tests: liver (median 0.330), lung (median 0.291), muscle (median 0.169), small intestine (median 0.240) and stomach (median 0.190). Brain had the largest median (0.405) but only 10 pairs and an interval crossing zero; kidney was near null (0.024).

Those medians sit on broad patient distributions (**Fig. 3b**). Each dot is one paired patient; the black bar marks the organ median and the number at right gives the fraction with a positive cosine.

Overall, 189 of 229 patients (82.5%) moved toward rather than away from fetal reference, although every organ contained heterogeneity. The bootstrap intervals in **Fig 3a** are conditional on the estimated developmental axes and do not propagate fetal- or adult-donor uncertainty.

I next asked whether a tumor fits its own organ’s fetal direction better than the alternatives (**Fig. 3c**). For each patient, the plotted value subtracts the median cosine to all mismatched organ axes from the cosine to the matched axis, using one identical protein set for every comparison. Positive values mean the matched fetal direction fits better, zero indicates no organ preference and negative values mean other organs fit better. The comparison retained a median of 2,252 proteins per primary-tier patient (range, 1,154-2,278); all exceeded the 500-protein floor. The organ-balanced matched advantage was 0.059, but its 95% organ-patient bootstrap interval crossed zero (-0.044 to 0.168). Liver, lung, small intestine and stomach showed FDR-significant positive matched-axis specificity. By contrast, kidney (median -0.055; 95% patient-bootstrap interval -0.080 to -0.020) and muscle (median -0.042; 95% patient-bootstrap interval -0.065 to -0.012) had negative matched advantages with intervals entirely below zero, whereas brain was inconclusive.

Direction does not tell us how far a tumor travels, so **Fig. 3d** reports a second quantity: the projection coefficient onto reverse maturation. A coefficient of 1 would reproduce the full fetal-to-adult gap in reverse; 0 would make no progress along that axis. The organ-balanced coefficient was 0.195 (95% organ-patient bootstrap interval, 0.069-0.244). Thus, even patients that point in the fetal direction typically traverse only part of the developmental distance. Within this dataset, the evidence supports partial alignment, not a wholesale return to fetal tissue.

### Fetal-like convergence overlaps with, but extends beyond, adult tissue-identity loss

**Figure 4a** shows five ways of scoring the same tumor-minus-adjacent change vectors. For each row, the point is the organ-balanced median cosine and the horizontal line is its 95% organ-patient bootstrap interval. Values above zero indicate that tumor change points in the named direction. The first row measures alignment with the complete reverse-maturation direction for that organ, from adult toward fetal tissue. The next two rows divide this developmental direction into a component shared across organs and a remaining organ-specific component, asking whether the fetal-like signal resembles a common developmental program or the fetal identity of the tissue of origin. Because these component directions are normalized separately, their cosines should not be added to reconstruct the full cosine. The final two rows examine alternative tumor programs after removing geometric overlap with directions already counted. “Identity loss after development” is the portion of movement away from the target adult-organ identity, toward other adult organs, that is not also reverse maturation. “Shared cancer after both” is the portion of the pan-cancer direction that is not aligned with either reverse maturation or identity loss. Thus, “after” describes the order used to remove overlap, not the biological order of events.

The complete reverse-maturation cosine was 0.240. Alignment with the cross-organ shared maturation component was nearly as strong at 0.219 (95% organ-patient bootstrap interval, 0.147-0.290), whereas alignment with the organ-specific component was smaller and less consistent at 0.139 (-0.021 to 0.243). Tumor change therefore aligned more clearly with a broadly shared fetal-like program than with the portion of fetal development unique to each organ. The unadjusted identity-loss cosine was 0.1225 but fell to 0.085 after its overlap with reverse maturation was removed, indicating that some apparent identity loss represented the same directional change as fetal-like convergence and that the remaining identity-specific signal was modest. In contrast, the shared-cancer cosine decreased only from 0.321 to 0.279 after removing overlap with both development and identity loss. A strong pan-cancer program therefore remained distinct from the fetal-like signal. These cosines measure directional similarity, not the fraction of total tumor remodeling explained by each program.

The apparent tension between becoming fetal-like and losing adult identity becomes explicit in **Fig. 4b**. For each organ, the dot is the cosine between the reverse maturation and identity loss axes before Gram-Schmidt orthogonalization; the angle label expresses the same geometry. A cosine of 1 would mean that the axes are identical, and 0 would mean that they are orthogonal. The primary-organ median was 0.524 (range, 0.293-0.705), corresponding to angles of roughly 73 to 45 degrees. Identity erosion is therefore part of the fetal direction, but substantial room remains for the two axes to differ. These fixed reference-axis values do not have patient bootstrap intervals.

Because a sequential decomposition assigns the overlap between reverse maturation and adult identity loss to whichever axis enters first, I repeated the analysis with their order reversed (**Fig. 4c**). Reverse maturation had an organ-balanced cosine of 0.240 when entered first and remained positive at 0.203 (95% organ-patient bootstrap interval, 0.129-0.239) after identity loss entered first. Adult identity loss had a cosine of 0.123 when entered first but decreased to 0.085 after reverse maturation entered first; the latter interval crossed zero (-0.137 to 0.202). Thus, fetal-like alignment was not explained solely by loss of adult identity: a positive reverse-maturation component remained after identity loss received first claim on their shared direction, whereas the remaining identity-loss component was smaller and uncertain after reverse maturation received first claim.

An order-invariant view reaches the same conclusion (**Fig. 4d**). The plane spanned jointly by development and identity captured 10.1% of paired change (95% organ-patient bootstrap interval, 6.9%-23.7%). The stacked bars use a Shapley allocation, which averages the credit assigned under both entry orders. Development received 60.2% of the joint space and identity the remaining 39.8%. This is a fair statistical division of shared explanatory space, not a causal partition. The label at the right of each bar gives the joint fraction of total paired change, so a large share should not be mistaken for a large fraction of the whole tumor response. The organ-balanced split also conceals heterogeneity: identity received the larger share in liver (60.0%) and kidney (76.0%), including liver despite its strong developmental alignment.

**Figure 3c** compared complete organ-level maturation vectors: it asked whether tumor change aligned more strongly with reverse maturation in the matched organ than with reverse maturation in other organs. **Figure 4e** asks a different question: within the matched organ’s vector, does the alignment come from maturation changes shared across organs or from changes specific to that organ? To answer this, I separated each fetal-to-adult maturation vector into a shared direction estimated from the other organs and a remaining organ-specific direction, then scored tumor-minus-adjacent change against the reverse of each. The median shared score was positive in every primary organ, whereas the organ-specific score ranged from strongly positive in brain to negative in muscle and kidney. Each line connects the two scores for the same organ. This helps reconcile the two panels: if much of the fetal-like alignment follows a direction shared across tissues, the matched organ’s complete vector need not align better than vectors from other organs.

The scale of the unexplained response is shown last (**Fig. 4f**). Each point is an organ median residual percentage with a patient-bootstrap interval, and the diamond is the organ-balanced estimate. A median 77.9% of paired proteomic change remained outside developmental reversion, adult identity loss and shared cancer directions. The candidate fetal-like signal is stable under these internal analyses, but it still describes only a minority of tumor remodeling.

### Suppression of adult-high proteins contributes most to fetal-like alignment

At the protein level, there are two ways to move toward fetal tissue. An adult-high protein can decrease in tumor, or a fetal-high protein can increase. To quantify these routes, I decomposed the dot product underlying the developmental-reversion cosine into protein-level contributions, multiplying each protein’s centered reverse-maturation value by its mean paired tumor change within each organ and retaining positive contributions. The stacked bars show the proportion of positive reversion contribution arising through each route (**Fig. 5a**), with an organ-balanced summary at the bottom. Suppression of adult-high proteins accounted for 65.5%, compared with 34.5% from fetal-high reactivation. Reactivation ranged from 25.3% in small intestine to 51.8% in brain across the primary organs.

**Fig. 5.**
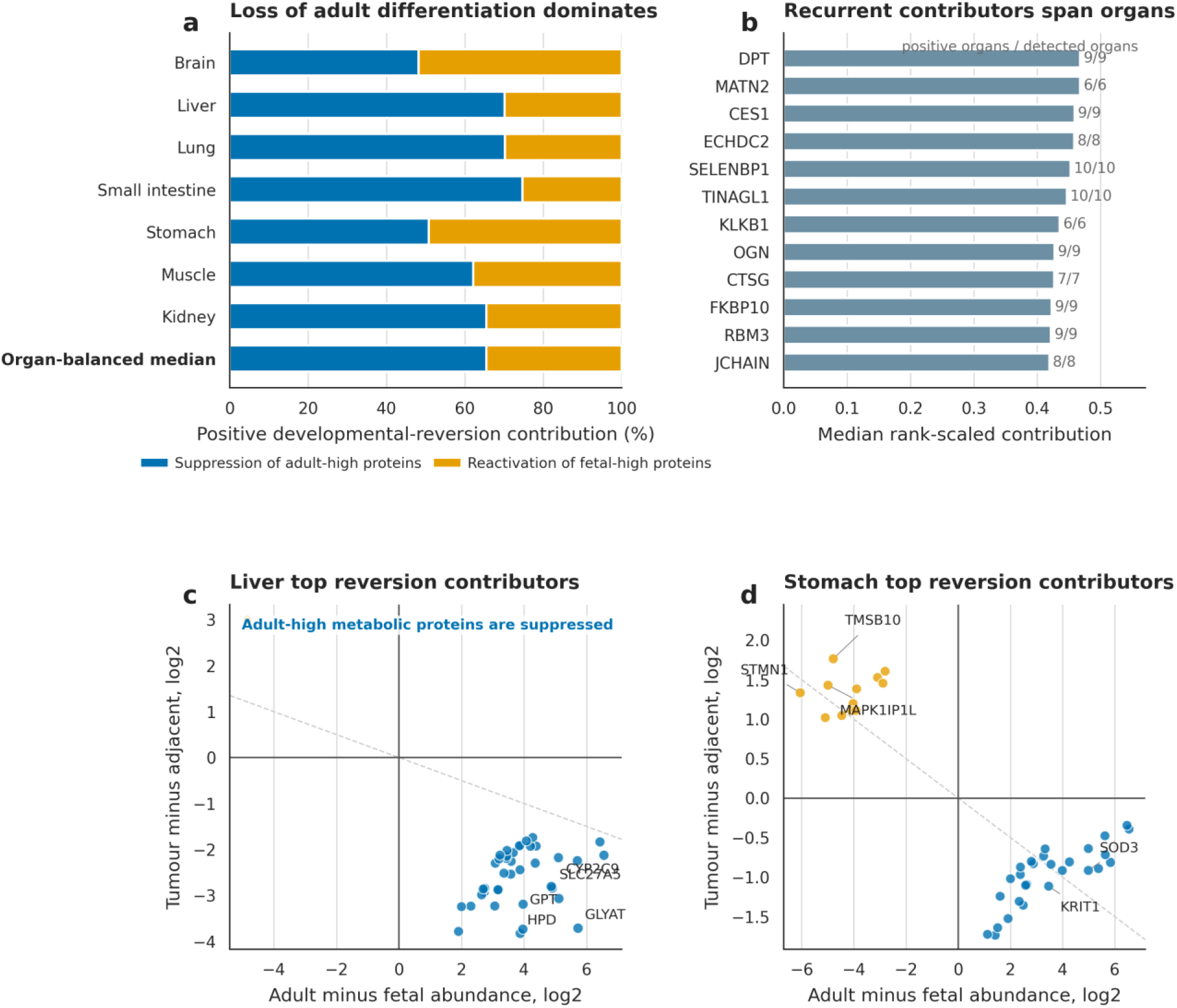
Fetal-like convergence receives most positive contribution from suppression of adult-high proteins. a, Share of positive developmental-reversion contribution arising from suppression of adult-high proteins or reactivation of fetal-high proteins. b, Top recurrent non-composition protein contributors after within-organ rank scaling; bar length summarizes relative rank across organs, and labels give positive organs over organs with sufficient detection. c, Top 40 positive liver contributors after excluding curated composition-program proteins. The x axis is adult minus fetal abundance, and the y axis is tumor minus adjacent abundance, so the lower-right quadrant represents suppression of adult-high proteins and the upper-left represents reactivation of fetal-high proteins. d, The corresponding stomach analysis. Labeled proteins had FDR-adjusted q<0.05 for paired tumor change.

The ranking in **Fig. 5b** asks which proteins contribute repeatedly across organs. Bar length is the median within-organ rank-scaled contribution, so it summarizes recurrence and relative importance rather than raw abundance; the labels report organs with a positive contribution among organs with adequate detection. DPT, MATN2, CES1, ECHDC2, SELENBP1 and TINAGL1 were the six highest-ranked recurrent contributors.

Liver makes the two molecular routes concrete (**Fig. 5c**). The x axis is adult minus fetal abundance, and the y axis is tumor minus matched adjacent abundance. The lower-right quadrant therefore contains proteins that are more abundant in adult than fetal liver but decrease in tumor relative to adjacent tissue. The upper-left quadrant represents the complementary route: proteins that are more abundant in fetal than adult liver and increase in tumor. Liver contributors were concentrated in the lower-right quadrant. The six highest-ranked were GLYAT, CYP2C9, HPD, GPT, SLC27A5 and AOX1, a candidate pattern consistent with tumor-associated loss of differentiated liver metabolic functions. The panel shows the top 40 positive contributors after excluding curated composition-program proteins; labels identify proteins with FDR-adjusted q<0.05 for paired tumor change.

Stomach uses the same coordinates and selection rule (**Fig. 5d**), but both positive-contribution quadrants are populated. Adult-high proteins decrease in tumor in the lower-right quadrant, whereas fetal-high TMSB10, STMN1 and MAPK1IP1L increase in the upper-left quadrant. These proteins contribute to the measured fetal-like transition but do not establish that any individual protein drives reprogramming. Some contributions may also reflect differences in stromal, immune or metabolic composition between tumor and adjacent tissue.

### Developmental alignment is stable across sensitivity analyses

The robustness summary begins with a heatmap of organ median developmental cosines (**Fig. 6a**). Organs are columns, and the main analysis plus five sensitivity variants are rows. Similar colors and numbers down a column mean that the organ estimate is stable to that analytical choice. Robust within-sample z-scoring and the cross-fitted adjacent reference preserved the sign in all 10 organs. Retaining intensity 1 as measured changed every organ median by at most 0.00005; brain changed from 0.4046 to 0.4045. Protein ranks were more sensitive: 35 of the liver top 40 overlapped between rules, so **Fig. 5** and the protein-level claims use the sentinel-excluded primary analysis. Removing 392 broadly defined immune, blood, matrix, stromal, endothelial, histone, mitochondrial and cell-cycle proteins preserved the sign in nine of 10 organs; only kidney shifted slightly below zero. Exact batch and instrument matching were possible for six organs (brain, kidney, lung, muscle, small intestine and stomach), and all remained positive. Blank cells indicate that no fetal and adult samples shared a technical stratum.

**Fig. 6.**
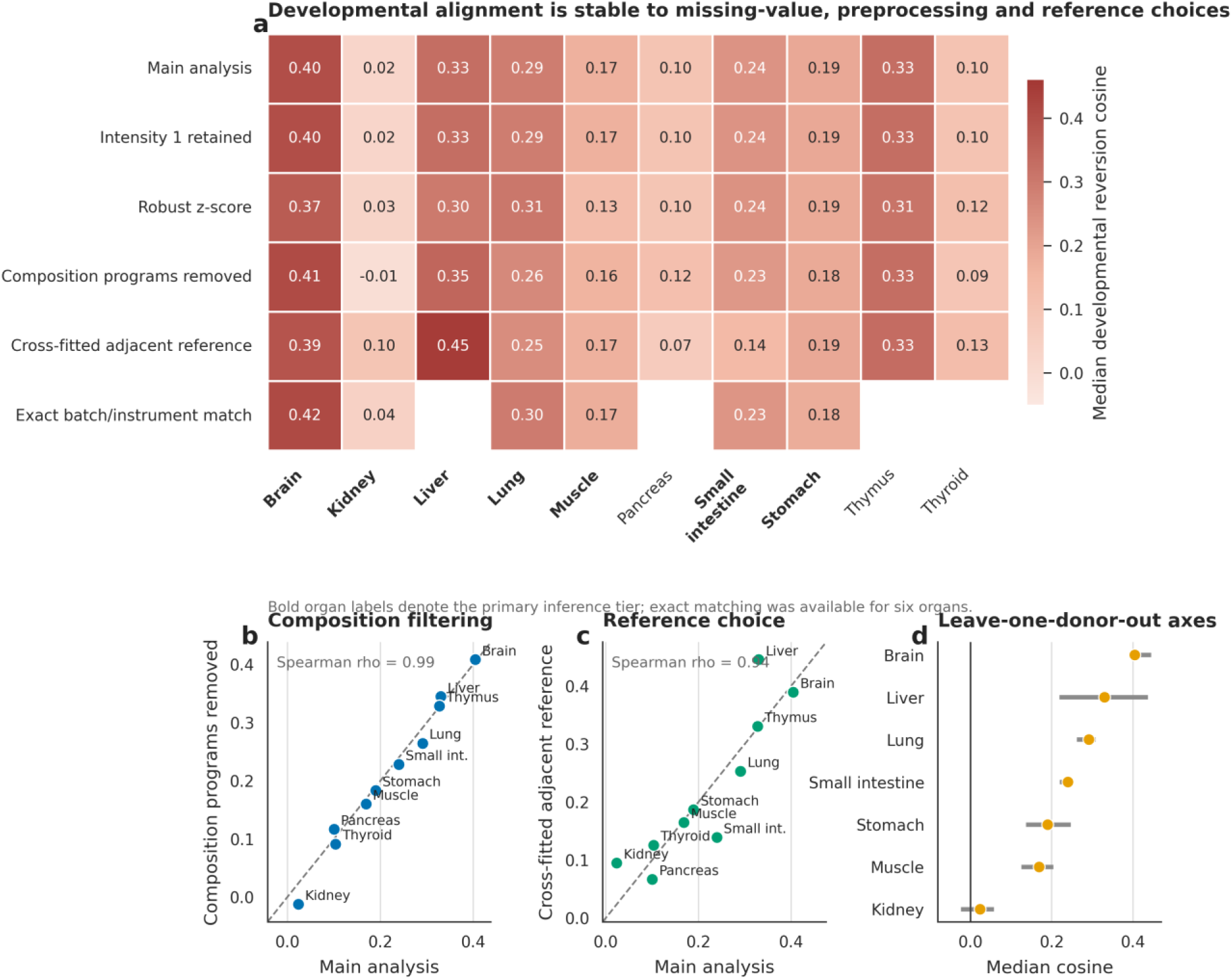
Developmental alignment is stable across missing-value, reference, scaling, composition and donor sensitivity analyses. a, Organ median developmental-reversion cosine, shown as values and color intensity, under the main analysis and five sensitivity analyses, including a comparator that retains exact-1 intensities as measured. Blank exact-match cells indicate that no fetal and adult samples shared an acquisition-batch and instrument stratum. Bold organ labels denote the primary tier. b, Main estimate on the x axis versus the estimate after removing curated composition programs on the y axis; the dashed line denotes equality. c, Main estimate versus a cross-fitted adjacent-normal developmental endpoint. d, Main organ median and full range after leaving out each fetal or adult donor in turn; ranges are stability summaries, not confidence intervals.

Composition filtering is unpacked in **Fig. 6b**. Each organ’s main estimate is plotted against its filtered estimate; points near the diagonal are unchanged; their Spearman correlation was 0.99, and kidney was the only organ to cross zero. This argues against the result being driven solely by the broad filtered programs but does not remove all cell-composition effects.

**Figure 6c** changes the reference rather than the protein set, replacing the external adult endpoint with a leave-one-patient-out adjacent-normal centroid from the same organ. The organ ranking remained similar (Spearman correlation, 0.94), showing that the directional result is not dependent on one adult reference cohort.

The final check omits one fetal or adult donor at a time (**Fig. 6d**). The orange point is the main organ median, and the grey segment is the full leave-one-donor-out range, not a confidence interval. Brain, liver, lung, muscle, small intestine and stomach remained positive across all omissions, whereas kidney crossed zero. These stress tests make a single preprocessing choice, broad composition program or measured batch stratum unlikely to explain the direction of the result, but they do not eliminate unmeasured preservation or cohort effects. The conclusion is therefore directional and partial; its magnitude and organ specificity should not be overinterpreted, especially where developmental donor counts are small.

## Discussion

This case study produced two linked conclusions. Biologically, tumor-minus-adjacent proteomic change in 229 patients across seven organs showed partial fetal-like convergence that overlapped with, but was not exhausted by, loss of adult tissue identity. Methodologically, a human-mediated, cross-model critique-and-refinement workflow converted a natural-language question into an inspectable, versioned re-analysis and exposed consequential defects along the way. The biological result remains a candidate discovery, and the workflow remains a single feasibility case rather than a reliability benchmark.

Conditional on the final audited pipeline, the biology supports a specific and limited model. Tumor change aligned with reverse maturation in most patients but typically traversed only part of the fetal-to-adult gap. Developmental and adult-identity-loss axes overlapped, yet a positive developmental component remained after identity loss entered first. Suppression of adult-high proteins supplied more of the positive alignment than reactivation of fetal-high proteins. At the same time, 77.9% of tumor-associated proteomic change remained outside development, identity loss and the shared-cancer axis, arguing against wholesale reversion to a fetal state.

The audit history calibrates confidence at different levels of the biological claim. The common-mask correction materially weakened matched-organ specificity, the missing-value correction barely moved aggregate geometry but changed part of the liver protein ranking, and the coupled-bootstrap correction changed uncertainty accounting without changing patient scores. A stable headline statistic did not guarantee a stable mechanistic ranking. Conversely, a detected implementation defect did not imply that every result was invalid. Versioned reruns made those distinctions observable rather than leaving them to intuition.

Human oversight was central even though it did not consist of manually writing or line-by-line certifying every code statement. The scientist selected the question, constrained the data and biology, proposed alternative explanations, interrogated figures, relayed and adjudicated critique, demanded reruns and accepted responsibility for interpretation. The primary model was treated as a code author rather than its own evaluator, generated prose was tied to derived tables, and corrections were preserved as named releases. This is a transparent division of labor, not a claim that conversational direction is equivalent to conventional programming expertise. For high-stakes conclusions, expert code review and independent reproduction remain stronger evidence.

Cross-model critique also has important limits. Agreement between two frontier model families is not statistical independence, and both systems may share training data, software conventions and reasoning biases. The critic was not blinded to the primary artifacts, and there was no manually written reference implementation, external benchmark or second analyst. The interaction record was reconstructed after the fact and does not include a literal export of the opening chat. This case therefore cannot estimate error rates, productivity gains or generalizability, and it does not establish audited vibe coding as a safe default.

Several biological limitations remain independent of code correctness. Fetal and adult tissues came from different donors rather than longitudinal developmental series. Developmental state is partly confounded with acquisition era, preservation and donor condition, and exact technical matching was available for only six organs. Bulk proteomics cannot distinguish malignant-cell reprogramming from changes in stromal, immune, vascular or parenchymal composition. Adjacent non-tumor tissue can contain field effects. Some developmental references contain only two donors, and the bootstrap intervals do not propagate developmental-donor uncertainty. The atlas lacks an external cohort with matched design for replication.

Together, these results support a calibrated dual conclusion. Partial fetal-like convergence is a biologically testable candidate that merits independent evaluation with spatial or single-cell proteomics and lineage-matched experimental models. Audited vibe coding is an inspectable way to conduct a targeted re-analysis when model-written code is treated as untrusted output, subjected to separate-model critique, corrected through versioned reruns and bound to executable checks. The biology and the workflow have different evidentiary limits, and neither should be judged by the fluency of the conversation that produced them.

## Methods

### Operational definition and study roles

For this case study, vibe coding was defined operationally as natural-language scientific direction with the primary large language model authoring essentially all executable analysis code. Audited vibe coding was defined as a human-mediated, cross-model critique-and-refinement workflow with four requirements: a separate model family critiqued the produced repository and claims; findings were returned to the primary conversation for correction; each correction produced a versioned rerun; and specified claims were checked against generated artifacts. Unlike same-model reflection or self-refinement,^20,21^ this case assigned the critic role to another model family, while following the broader logic of model-based critique.^22^ The human scientist remained responsible for question selection, biological constraints, interpretation, audit adjudication and final approval. The human scientist did not claim to have manually authored or line-by-line verified every implementation detail.

### Interaction and input reconstruction

The workflow was reconstructed from the versioned reproducibility packages, timestamps, source-code constants, generated READMEs, audit-correction logs, claim manifests and release-verification reports. The executable pipeline reads four source workbooks: Protein intensity matrix.xlsx, Adult normal tissue matrix.xlsx, Paired tumor-adjacent tissue matrix.xlsx and Sample origin information.xlsx. The literal initial prompt sequence was not exported, so exact prompt wording, turn counts and token counts are not reported.

### Primary model workflow and human steering

The primary environment was the ChatGPT web interface in early August 2026 using GPT-5.6 Sol with Max reasoning. The interaction was iterative rather than a single master prompt. The model inspected data dimensions and labels, proposed a vector-based analysis, wrote and executed Python, created derived tables and figures, and drafted the manuscript. The human scientist reviewed biological plausibility and communication, asked targeted questions about figure relationships and mathematical interpretation, requested alternative analyses such as the identity-loss order test, and supplied audit findings for correction.

### Cross-model critique and versioning

Audits used Opus 5 within Claude Code version 2.1.226. The auditor was instructed to compare the paper, code, derived data and figures, with attention to cohort construction, masks, normalization, missing values, vector definitions, uncertainty, multiple testing, claim binding and portability. Written findings were pasted into the primary ChatGPT thread. The primary model implemented changes, reran the analysis where required and generated a new named package. Version 2 documented 12 findings; version 3 listed eight follow-up findings; version 6 documented eight ingestion and provenance findings; and version 7 documented four methods-consistency findings. Version 4 was a human-directed interpretability revision, and version 5 was a human-directed scientific extension.

### Release verification

Release checks included deterministic random seeds; source and converted-file SHA-256 hashes, sizes and dimensions; raw-value and missing-sentinel counts; cache rejection on provenance mismatch; consistency assertions between manuscript claims and CSV or JSON outputs; patient-row and organ-summary equality across versions when invariance was expected; equality of the two development-identity subspaces across entry orders; Shapley-share accounting; exact complementarity of coupled intervals; multiplicity classifications; workbook formula scans; source-figure hashes; accessibility checks; and rendered inspection of manuscript, figure-set and workbook pages. These tests were selected during the iterative audit process and therefore should be interpreted as regression and consistency checks, not as a prospectively complete test suite.

### Data source and analysis cohorts

I analyzed the protein-intensity, adult-normal, paired tumor-adjacent and sample-origin workbooks released with the TPHP human proteome atlas.^9^ The workbooks provided were: Sample origin information.xlsx, Protein intensity matrix.xlsx, Adult normal tissue matrix.xlsx and Paired tumor-adjacent tissue matrix.xlsx. The full protein matrix contained 2,856 samples and 13,609 protein groups. The adult-normal matrix contained 466 samples and 10,170 proteins. The paired matrix contained 1,780 rows representing 890 complete tumors and adjacent non-tumor patient pairs, with 13,207 proteins. Analyses were restricted to the 9,928 proteins common to all three matrices. Fetal samples flagged for low protein identifications were excluded.

Fetal labels were mapped before analysis using a prespecified, explicit organ dictionary. Fetal diencephalon, mesencephalon, metencephalon, myelencephalon and telencephalon were pooled as brain; fetal kidney and metanephros were pooled as kidney; and fetal intestine and small intestine were pooled as small intestine. Fetal liver, lung, muscle, pancreas, stomach, thymus gland and thyroid gland mapped directly to the corresponding adult organ. The remaining 15 fetal tissue labels were excluded because they lacked a prespecified unambiguous fetal, adult and cancer-organ match. Fetal gonad was not mapped to adult testis or testicular germ-cell cancer because the fetal label did not specify sex or lineage.

Replicate acquisitions from the same fetal donor and organ, adult donor and organ, or cancer patient, organ and state were collapsed by the median. The primary evidence tier required at least two fetal donors, two adult donors and paired cancer samples for an organ. Brain, kidney, liver, lung, muscle, small intestine and stomach met this criterion. Pancreas, thymus and thyroid were analyzed only in sensitivity summaries because one developmental endpoint had a single donor.

### Normalization and protein eligibility

Stored intensity values at or below 1 were treated as missing. Within the 9,928-protein common universe, exact 1 occurred 12,165 times in the atlas matrix and 7,641 times in the paired matrix. The corresponding source-wide counts were 26,700 and 17,957. Exact 1 was absent from the adult-normal matrix and was separated from the next positive stored intensity (12.77). This discrete floor was therefore treated as a missing-value sentinel. Intensities were log2-transformed and median-centered within each sample; missing values were not imputed. Source workbooks were streamed from stored OOXML values rather than displayed cell text, and cached conversions were reused only when source and CSV hashes matched. Within each directly matched organ, a protein was eligible if detected in at least half of fetal biological units, at least half of adult biological units and at least half of paired cancer changes. Patient scores required at least 500 jointly finite proteins across the cancer vector and all reference axes; the median number used ranged from 2,832 to 5,322 across primary organs. Muscle, with two fetal and two adult donors, had the lower value.

### Developmental, identity and shared cancer axes

For organ o, the maturation vector D(o) was the protein-wise median adult profile minus the median fetal profile. Developmental reversion was -D(o). The adult identity vector I(o) was the adult organ centroid minus the protein-wise median centroid of all other adult organs; identity loss was -I(o). For each cancer organ, paired change vectors were calculated as tumor minus adjacent non-tumor abundance. Their organ-level centroids were median-centered and normalized to unit length. The shared cancer vector U(-o) was the mean of these normalized centroids across all other cancer organs, so the organ being tested did not contribute to its own reference.

For every patient, the cancer-change vector and reference axes were median-centered over jointly eligible proteins and scaled to unit length. The identity-loss axis was orthogonalized against reverse maturation. The shared cancer axis was then orthogonalized against both preceding axes. Cosine similarities were dot products between the patient’s unit cancer-change vector and each orthonormal axis. The reverse-development projection coefficient was the scalar projection of cancer change onto reverse maturation divided by the squared length of the maturation vector; a value of 1 corresponds to traversing the complete fetal-to-adult reference gap in reverse. Signed fractions were sign(cosine) multiplied by cosine squared. The residual fraction was one minus the sum of the three unsigned squared cosines.

I assessed developmental-identity overlap in three complementary ways. First, the cosine and angle between reverse maturation and identity loss were calculated on each organ’s fixed eligibility set. These reference-axis values were not assigned patient-bootstrap intervals. Second, the sequential decomposition was repeated with identity entered first and development orthogonalized against it. Third, the squared projection of each unit cancer vector onto the two-axis plane was calculated under both orthonormal bases and averaged; the maximum numerical discrepancy between entry orders was 2.00e-15. This joint fraction is order-invariant.

Credit within the joint developmental-identity subspace was allocated with a two-variable Shapley calculation. For each patient and each axis, I averaged its squared cosine when entered alone and its incremental squared projection when entered second. The two allocations sum to the joint subspace fraction. Dividing each by that joint fraction gives the displayed shares. Development and identity share intervals were estimated together from the same resampled patients and organs; because the two shares sum to 1 in every replicate, their percentile intervals are exact complements. This procedure provides an order-averaged statistical allocation and does not identify causal mediation.

Matched-axis specificity was analysed separately from the orthogonal decomposition. Within each patient, the matched developmental cosine and every mismatched-organ developmental cosine were computed on exactly the same proteins: the target organ’s eligibility mask intersected with proteins having finite maturation estimates for all compared organs. The matched advantage was the matched cosine minus the median mismatched cosine. Among primary-tier patients, this comparison used a median of 2,252 proteins per patient (range, 1,154-2,278); every patient retained at least 500 proteins.

### Shared and organ-specific maturation

To divide maturation itself, each direct organ was excluded in turn and a shared maturation direction was estimated as the mean of unit-normalized maturation vectors from the other directly matched organs. The target organ’s maturation vector was regressed on this shared direction; the orthogonal residual defined organ-specific maturation. Patient cancer-change vectors were scored against the negative shared and organ-specific components separately.

### Statistical inference

Organ-level effects are patient medians, with 95% patient-bootstrap intervals estimated from 4,000 resamples within each organ. One-sided sign-flip tests used 20,000 random sign assignments to the patient scores, with a +1 correction, and were adjusted across 10 direct organs using the Benjamini-Hochberg procedure. Headline 95% organ-patient bootstrap intervals used an organ-balanced hierarchical bootstrap with 6,000 iterations: organs were sampled with replacement, patients were resampled within each selected organ, and the median of organ medians was retained. This prevents large cohorts from dominating the pan-organ estimate. These are organ-patient bootstrap intervals conditional on estimated maturation axes. Fetal and adult donors were not resampled, so the intervals do not propagate developmental-axis uncertainty; leave-one-donor-out ranges were reported separately as stability analyses, not confidence intervals.

### Sensitivity analyses

Four prespecified sensitivity analyses were performed. First, log2 median-centered values were divided by 1.4826 times the within-sample median absolute deviation about zero; non-finite scale estimates and scales below 0.05 were replaced with 1.0. Second, 392 proteins were removed using exact markers and the prefixes IG, HLA, TRAV, TRBV, TRDV, TRGV, FCGR, FCER, COL, LAMA, LAMB, LAMC, MMP, TIMP, ITGA, ITGB, MCM, CCNA, CCNB, CDK, CENP, KIF, HIST and NDUF. This intentionally broad set spans immune and immunoglobulin-related proteins, including IGF, IGFBP and IGSF matches; blood, extracellular-matrix, stromal and endothelial markers; histones; mitochondrial proteins; and cell-cycle programs, including CDKN matches. Third, fetal and adult maturation vectors were recomputed within exactly overlapping acquisition-batch and instrument strata where available, with strata weighted by the geometric mean of fetal and adult donor counts. Fourth, the adult endpoint was replaced with a leave-one-patient-out centroid of adjacent non-tumor samples from the same organ, preventing a patient’s adjacent sample from contributing to its own reference. Maturation-axis stability was also assessed by leaving out each fetal or adult donor in turn. A targeted fifth sensitivity retained exact-1 intensities as measured, using the same raw-precision conversion, cohort, seeds and scoring code. This isolates the sentinel rule from conversion precision and all other analytical choices.

### Protein contribution analysis

For each organ, I represented reverse maturation and mean paired tumor change as protein-dimensional vectors, with each protein defining one coordinate. Both vectors were median-centered across their protein coordinates. I then decomposed their dot product into protein-level terms. For protein (i), the developmental-reversion contribution was the product of its centered reverse-maturation coordinate and its centered mean tumor-change coordinate. A positive product indicates that the protein changed in the reverse-maturation direction. Positive contributions were classified as suppression of an adult-high protein when the protein was more abundant in adult than fetal tissue and decreased in tumor, or as reactivation of a fetal-high protein when it was more abundant in fetal than adult tissue and increased in tumor.

For each protein, tumor change was evaluated by a one-sample t-test of the patient-level paired tumor-minus-adjacent values against zero, followed by Benjamini-Hochberg correction. To identify recurrent contributors across organs, contributions were rank-scaled within each organ and averaged across organs with sufficient protein detection. For the liver and stomach display panels, curated composition-program proteins were excluded before selecting the 40 largest positive contributions.

### Software, reproducibility and generative AI

Analyses were implemented in Python using pandas, NumPy, SciPy, matplotlib and seaborn. The random seed was 10,660. Complete analysis code is available from GitHub (https://github.com/xomicsdatascience/cancer-fetal-reversion). Conversion manifests record source and CSV SHA-256 hashes, sizes, dimensions, fractional-value checks and converter version. GPT-5.6 Sol in the ChatGPT web interface generated the analytical code, visualization code and manuscript drafts under human direction. Opus 5 in Claude Code version 2.1.226 performed separate repository and claim audits. Audit findings were relayed to the primary conversation for correction. No claim is made that all possible errors were found; the version logs document only identified findings and specified verification checks.

## Data availability

The source protein matrices and metadata were released with the TPHP atlas and are described in the original publication.^9^ The accompanying repository contains only derived results and figures; access to source data should cite and follow the terms of the original resource (https://db.prottalks.com/download.html).

## Code and interaction materials availability

The analysis script, derived patient and organ tables, sensitivity summaries, protein-driver tables, figure-generation code, versioned audit-correction logs, claim manifests and verification reports are available at https://github.com/xomicsdatascience/cancer-fetal-reversion. The literal opening chat transcript was not exported and is not represented as part of the reproducibility record.

## Author contributions and AI assistance

J.G.M. supplied the data context, directed the analyses, reviewed outputs, proposed alternative explanations, adjudicated audit findings, interpreted the biology and approved the manuscript. ChatGPT GPT-5.6 Sol generated analysis code, executed analyses, created figures and drafted text under those instructions. Opus 5 in Claude Code audited the code, outputs and claims. The AI systems are not authors.

